# An Integrated Platform for in vivo Electrophysiology in Spatial Cognition Experiments

**DOI:** 10.1101/2023.08.02.551691

**Authors:** A. Brea Guerrero, M. Oijala, S. C. Moseley, T. Tang, F. Fletcher, Y. Zheng, L.M. Sanchez, B. J. Clark, B. L. Mcnaughton, A. A. Wilber

**Affiliations:** Florida State University, Program in Neuroscience, Tallahassee, FL; Psychology Department, Florida State University, Tallahassee, FL; Univ. of California, Irvine, Irvine, CA; Dept. of Psychology, The University of New Mexico, Albuquerque, NM; The Univ. of Lethbridge, Lethbridge, AB, Canada

## Abstract

Spatial cognition research requires behavioral paradigms that can distinguish between different navigational elements, such as allocentric (map-like) navigation and egocentric (e.g., body centered) navigation. To fill this need, we developed a flexible experimental platform that can be quickly modified without the need for significant changes to software and hardware. In this paper, we present this inexpensive and flexible behavioral platform paired with software which we are making freely available.

Our behavioral platform serves as the foundation for a range of experiments, and though developed for assessing spatial cognition, it also has applications in the non-spatial domain of behavioral testing. There are two components of the software platform, ‘Maze’ and ‘Stim Trigger’. Both programs can work in conjunction with electrophysiology acquisition systems, allowing for precise time stamping of neural events with behavior. The Maze program includes functionality for automatic reward delivery based on user defined zones. ‘Stim Trigger’ permits control of brain stimulation via any equipment that can be paired with an Arduino board. We seek to share our software and leverage the potential by expanding functionality in the future to meet the needs of a larger community of researchers.

**Significance Statement:** This paper presents an innovative and cost-effective behavioral platform designed to distinguish between different navigational elements, addressing the crucial need for better spatial cognition research paradigms. The platform’s flexibility allows for quick modifications without major software or hardware changes. Additionally, the freely available software, comprising ‘Maze’ and ‘Stim Trigger’ components, enables precise time stamping of neural events with behavior, while facilitating automatic reward delivery and brain stimulation control. Beyond spatial cognition assessment, the platform’s adaptability extends to non-spatial behavioral testing. By openly sharing this software, the authors aim to foster collaboration and encourage future developments, promoting its application to a broader community of researchers. This platform represents a significant advancement in spatial cognition research and behavioral experimentation methods.

## Introduction

Spatial cognition is a burgeoning research field in neuroscience that has implications in memory, aging, neurodevelopmental disorders, and progressive neurologic disorders. Spatial memory is present in most animal species (Benhamou and Poucet, 1995), and it is a key element in how individuals interact with their surroundings. The use of animal models is common because they provide methodological benefits, such as the ability to utilize invasive electrophysiological recording or imaging devices in pertinent brain regions.

The use of animal models in research requires the creation of innovative tools that enable scientists to investigate new concepts related to spatial navigation, learning and memory, anxiety like behavior, decision making and other behavioral variables of interest. Researchers commonly employ mazes to assess spatial cognition, learning, and memory (Paul et al., 2009). These include the radial arm maze, T and Y mazes, Morris water maze, and Barnes maze, which are among the most frequently used (Gawel et al., 2018). The design of each maze and the task’s protocol dictate which aspects of spatial memory and cognition are evaluated.

The Morris water maze and Barnes maze are examples of open mazes that offer multiple paths to navigate towards a designated objective, typically a shelter or escape platform, using the cues surrounding the testing area (Morris, 1981; Barnes, 1988). The task-learning process generally relies on negative reinforcers; however, the stress induction is much stronger in the case of the Morris water maze, as measured in plasma corticosterone (Harrison et al., 2009). Alternatively, the radial arm maze, the T and Y mazes, as well as versions such as the Cincinnati maze offer restricted route options (Paul et al., 2009). These mazes have led to significant advances in the field, for example place cells were first observed in a restricted-route maze (O’Keefe, 1976). Restricted route mazes are often used to study working memory and typically use positive reinforcers like food and water to motivate behavior(Dudchenko, 2004; Morellini, 2013). Albeit with some exceptions, the aforementioned mazes focus on allocentric (map-like) processing of space. Allocentric processing relies on distal cues or ‘landmarks’ to create a spatial reference frame (Bermudez-Contreras et al., 2020). A remarkable exception is the study published by Rondi-Reig and collaborators (2006), showing that their ‘starmaze’ could be used to develop allocentric, and sequential-egocentric tasks (Rondi-Reig et al., 2006). Furthermore, studies on animal models, such as rodents navigating mazes, have shown that they rely more on local cues and directional signals to guide their movements, which is consistent with stimulus-response learning rather than true egocentric navigation based on self-positioning within the environment (TOLMAN, 1948; Geerts et al., 2020). Thus, there is a need for experimental designs that further investigate the egocentric (e.g., body-centered) frame as well as the interaction between both reference settings.

We developed this platform to fill a need for a flexible platform to advance spatial cognition research using novel experimental designs and apparatuses including those that go beyond allocentric spatial location processing. It is common practice to modify mazes and protocols, as seen in combinations like a radial arm maze submerged in a water tank (Buresová et al., 1985). That is why we created a cost-efficient maze with a design based on a circular platform, similar to those used in the Barnes Maze, which, when combined with custom MATLAB software, enables researchers to quickly develop customized experimental designs both within the field of spatial cognition research and beyond.

Recording the activity of neurons in freely moving animals during task performance provides valuable insight into how different areas of the brain communicate with each other. The brain dynamics between areas such as the hippocampus, entorhinal cortex, subiculum, thalamus, retrosplenial, and parietal cortex are known to be critical for spatial cognition and other high-level cognitive processes like learning, memory, decision making, and attention (Bermudez-Contreras et al., 2020). To record neuronal activity simultaneously, researchers use recording arrays with tetrodes or multichannel silicon probes. Additionally, silicone probes offer the ability to record interactions within layered structures more easily, which is essential for understanding brain computations (Buzsáki et al., 2003).

Since the first publication using this software platform with tetrodes (Bower et al., 2005), we have developed and integrated the software to facilitate *in vivo* electrophysiological experiments using several recording platforms. The tasks that have employed it so far are focused on spatial cognition; however, the platform is flexible enough to allow for testing using many paradigms such as decision making. To achieve our goal, we implemented two commonly used electrophysiology recording platforms, Neuralynx and Open-Ephys, along with two real time video tracking approaches. We chose these platforms because they offer complementary features. Neuralynx is widely used for its excellent analog signal processing capabilities (though newer systems are capable of analog and digital processing), while OpenEphys provides a cost-effective and open-source solution for digital signal processing. Accurate interpretation of the behavioral correlates of brain signals depends on precise synchronization of video and brain signals. Neuralynx includes the necessary video tracking features, while Open-Ephys can be paired other platforms such as Bonsai, an open-source software program for real-time video analysis and more. Furthermore, our platform has built in functionality for optogenetic experiments, which can modulate brain activity with high specificity using TTL signals to trigger laser pulses at specific user defined points in the maze.

Our platform provides the hardware and software components necessary to flexibly and precisely measure brain signals during custom-designed tasks in freely moving animals if dictated by the experimental design or to conduct behavioral experiments without recording brain activity. Additionally, it can be combined with *in vivo* brain imaging techniques, like the UCLA Miniscope. We aim to collaborate to offer great flexibility in experimental design. cognitive processes.

## Materials and Methods

### Hardware

#### Platform

The circular platform (T-60RT, Unfinished Furniture of Wilmington) is a 60-inch diameter Parawood table top, 3/4 inch thick, and with a reverse bevel edge. To seal the surface and improve paint adhesion we used KILZ® water-based acrylic primer. The paint used was Chalky Finish Krylon “Misty Gray”. The color was selected to provide a neutral background to offer contrast with the rodents’ coat color, which in our case is white, brown or black for video tracking purposes. However, other colors can be used to increase this contrast for other combinations of rodent strains or to mimic the color of the proximal or distal walls.

The wall is made of low-density polyethylene and has a dual function: it holds the electronic components at a proper height for the animals to interact with them, and serves as a boundary for the animals, helping them use distal cues and avoid unintentionally crossing the edge of the maze. Different wall heights and other surface shapes can be used to assess the role of boundaries in the hippocampal processing of space, as observed in the activity of ‘boundary cells’ (Hartley et al., 2000; Solstad et al., 2008). Our software and hardware platform can work with any maze shape or apparatus. Moreover, wall color and height can be manipulated to investigate the impact of its use as a proximal cue. For example, by matching the color of the floor, the effect of walls as a proximal cue can be further reduced. Furthermore, the walls can be removed if the platform is placed high enough to reduce the chances of an animal jumping off (Gawel et al., 2018).

#### LED Cue Lights

We installed 32 evenly spaced CHANZON© LED lights (3V, 20mA) on the maze wall using eBoot© JST SM 2 Pin Plugs. In our configuration, each diode is connected to a 1.1K resistor to reduce its brightness. Different resistors can be used to adjust the LEDs to the desired brightness.

The wiring of electronic components, including the LEDs, uses XHF lever connectors for safety and easy modification. The LEDs are connected to a 48-channel USB-DIO-48 module through a Sysly(c) IDC50-B breakout board **(Fig 1).**

**Figure 1:**
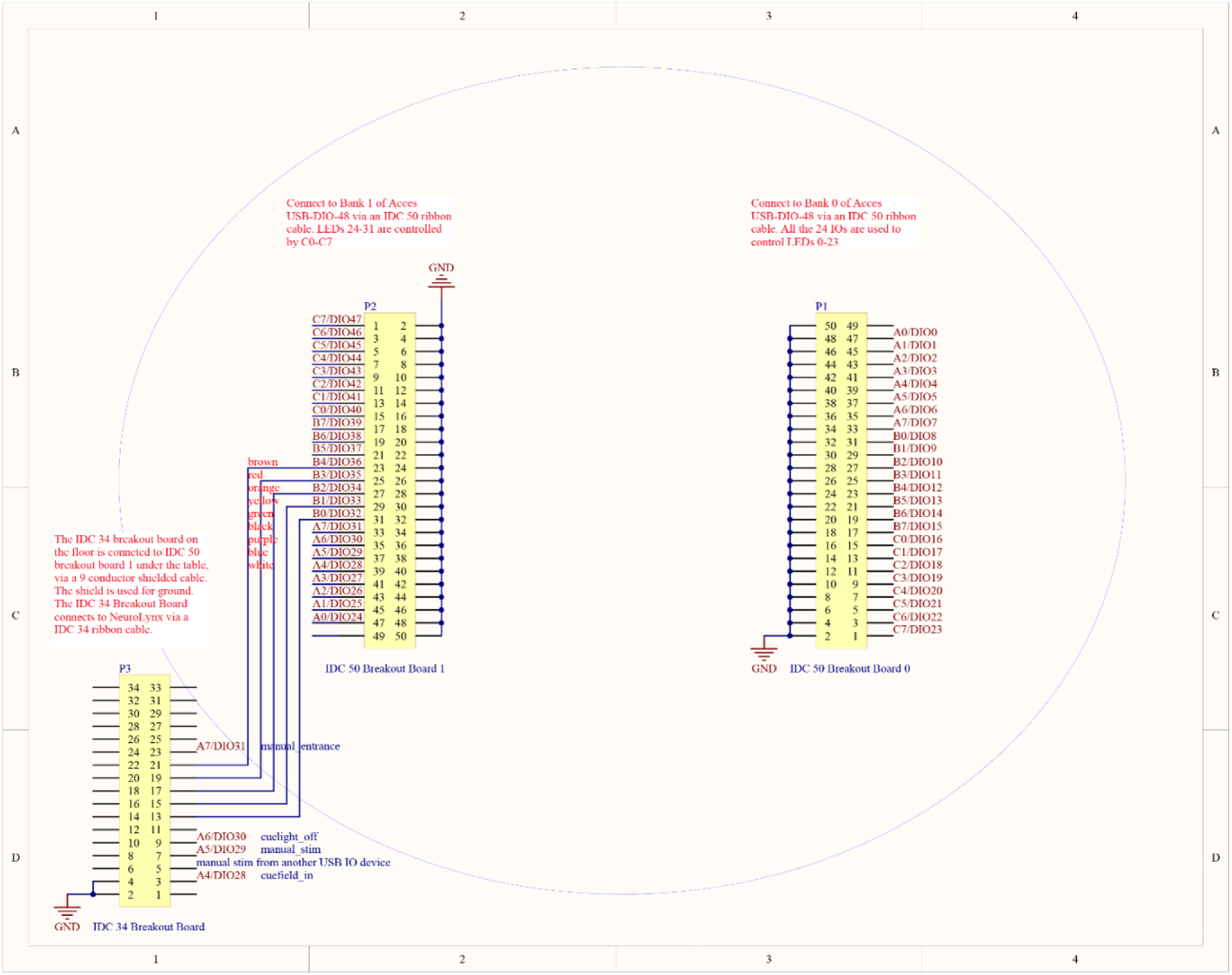
Pinout mapping for IDC Breakout Boards in the behavioral platform connecting the electronic components to USB Access I/O modules. Each pin offers a connection to control LEDs, solenoids, or send TTL signals.

Our Maze software (MATLAB(c)) controls 48 TTL-compatible bits in the I/O module using the AIOUSB API library (Anon, n.d.). The LED lights can be used as cues during experimental procedures and to automatically calibrate the maze location and zones of interest in the camera frame.

### Hardware systems

#### Liquid reward

To deliver liquid rewards to the animals, we installed eight solenoid-operated pinch valves (Cole-Parmer©) that are connected to tubes containing the liquid. The valves allow for delivery of water or nutrition drinks (Ensure®) to animals undergoing water or food restriction, respectively. The tubes are connected to evenly spaced spouts on the maze walls, with one spout placed over every fourth LED. Using less spouts would not require any modification to the maze software.

These valves are controlled using a 16-channel Digital I/O USB Module (USB-IDO-16, (Anon, n.d.). The Maze software (MATLAB©) uses the aforementioned AIOUSB API library (Anon, n.d.)} to control the delivery of the liquid.

#### Brain stimulation

Stimulation of the medial forebrain bundle is a strong reinforcer that is extensively used in animal models (OLDS and MILNER, 1954; Wise and Rompre, 1989; Wise, 1996; Wise, 2002). Various laboratories have successfully used brain stimulation as a reinforcer in spatial cognition tasks in both mice and rats (Euston et al., 2007; Wilber et al., 2014; Wilber et al., 2017; Stimmell et al., 2019; Cushing et al., 2020). Further, optogenetics is a powerful tool for modulating brain activity with temporal and spatial specificity (Emiliani et al., 2022).

We provide two options for signal to trigger stimulation, unipolar or bipolar. We use a TTL signal designed to trigger a bipolar output by a constant current stimulus isolator (SYS-A365 World Precision Instruments©). A data acquisition device (USB-1208FS-Plus, Measurement Computing) gates the TTL pulses used to control stimulation parameters via MATLAB using the MATLAB© Data Acquisition Toolbox. The characteristics of the stimulation, such as frequency, duration and duty cycle can be determined in two separate software packages: ‘Stim Trigger’ (for testing stimulation parameters alone in an operant chamber) and Maze (for delivering brain stimulation during experiments on the maze platform (MATLAB© Runtime 9.2). For further details on selecting stimulation parameters for rodents see Carlezon and Chartoff, 2007. Duty cycle determines the percentage of each stimulus cycle that is set as positive however the number of stimuli depends on the duration and frequency.

The unipolar TTL signal from the data acquisition device can be used as input for optogenetic laser control. The device can connect to the power supply for the laser via BNC connector. The signal features can be determined in our software similarly to those for electrical stimulation.

### Electrophysiology data acquisition

#### Headstages and Probes

We have used a variety of configurations for silicon probes. However, we will focus on the mice configuration using a single, H5 probe (Cambridge Neurotech). We attached the probe to an implantable nano-Drive. The probe was connected to a mini-Amplifier *\*emph{via} an interposer board (Cambridge Neurotech). We used a custom-made cable (Neuralynx©), containing SPI and MDR50 male connectors to connect the amplifier to a commutator (Saturn, Neuralynx©) which helped to reduce torsion from the freely moving animal and safeguard the integrity of the components. We used a second custom-cable to connect the commutator to either the OpenEphys Acquisition Board or the Neuralynx© Digital Lynx SX acquisition system.

For tetrode recording, we used recording arrays built in-house that consisted of 18 independently drivable tetrodes (Kloosterman et al., 2009; Wilber et al., 2017). We have used a variety of electrode interface boards to interface with Neuralynx© Digital Lynx SX: EIB-72-QC-Small for mice, and EIB-72-QC-Large, EIB-27-18TT, EIB-36-16TT, HS-72-QC-LED for rat recordings (Neuralynx©).

#### Acquisition systems

The Open Ephys© Acquisition Board and the Digital Lynx SX by Neuralynx© are two popular systems for acquiring extracellular electrophysiology data. The Open Ephys board offers an interface between up to 512 channels of data and the computer via USB connection. As the project is open source, components and assembly instructions are freely available. Alternatively, pre-assembled components and remote training can be purchased (Open Ephys and Contributors, 2022).

The Digital Lynx SX offers up to 512 channels with a modular configuration that can be customized to meet the needs and budget of experimenters. Additionally, it includes a Hardware Processing Platform for real-time closed loop data processing, and other advanced experiment designs using platforms such as MATLAB© (Neuralynx, 2022).

Both systems offer a range of features and flexibility, making them popular choices for researchers conducting electrophysiology experiments.

### Software

Our software is available in an OSF repository at: https://osf.io/svtzr/

Animal behavioral tracking can be accomplished using a range of software options. EthoVision is a frequently used video tracking software that captures animal movements in real-time and provides detailed behavioral data. EthoVisionXT extends the capabilities of EthoVision by offering a ‘Trial and hardware control’ module, which requires pairing with their USB-IO box. Noldus provides a complete hardware and software catalog for behavioral studies with rodents, but the modular nature of their software can potentially result in limitations if all required modules are not accessible, and their proprietary software lacks compatibility with other platforms such as Python or MATLAB.

ANY-maze is a versatile software that offers a wide range of capabilities for tracking animal movement and behavior, including support for video tracking and operant conditioning chambers. Importantly, it does not have ability to synchronize with other software platforms such as Python or MATLAB, nonetheless it is possible for users to code plug-ins to acomplish these functions. ANY-maze’s full license provides unlimited access to updates as well as add-ons. They also offer a range of ANY-maze interface devices for controlling hardware.

Bonsai-RX is a popular open-source software platform for real-time behavioral analysis and closed-loop experiments in neuroscience research. It provides a flexible environment for experimental design, allowing users to create, modify, and execute complex behavioral experiments with ease. Its extensive library of pre-built components and user-friendly interface simplifies the process of creating custom experiments that incorporate multiple data streams. Bonsai-RX is highly adaptable and its modular design enables researchers to incorporate a range of hardware and software components into their experiments, and this is the reason why we chose to implement it with our platform.

DeepLabCut is a widely used open-source software tool that uses deep learning algorithms to accurately and automatically track the position of key points on the subject’s body. When paired with Bonsai-RX, DeepLabCut provides a powerful toolset for closed-loop experiments and real-time behavioral analysis. Researchers can create complex experimental setups that incorporate multiple data streams, such as video tracking, electrophysiology, and optogenetics, and perform real-time analysis and closed-loop experiments. As it has already synergic functions with Bonsai-RX users could easily implement it with out software,

Researchers can transfer animal location data to MATLAB using UDP connections in Bonsai-RX or DeepLabCut. MATLAB can receive the data and process it for further analysis, visualization, or hardware control. The compatibility of these software solutions with other platforms provides researchers with the freedom to pair them with many other software solutions.

#### Maze

In developing our maze software, we utilized the App Designer platform, a visual environment in MATLAB that enables the rapid creation of graphical user interfaces. This choice provides users with the ability to modify our code package in a relatively straightforward manner to better suit their needs. Some examples of the use of this software with further explanation of the functionalities used can be found in the Results section. The GUI includes the following functionalities **(Fig 2)**.

**Figure 2:**
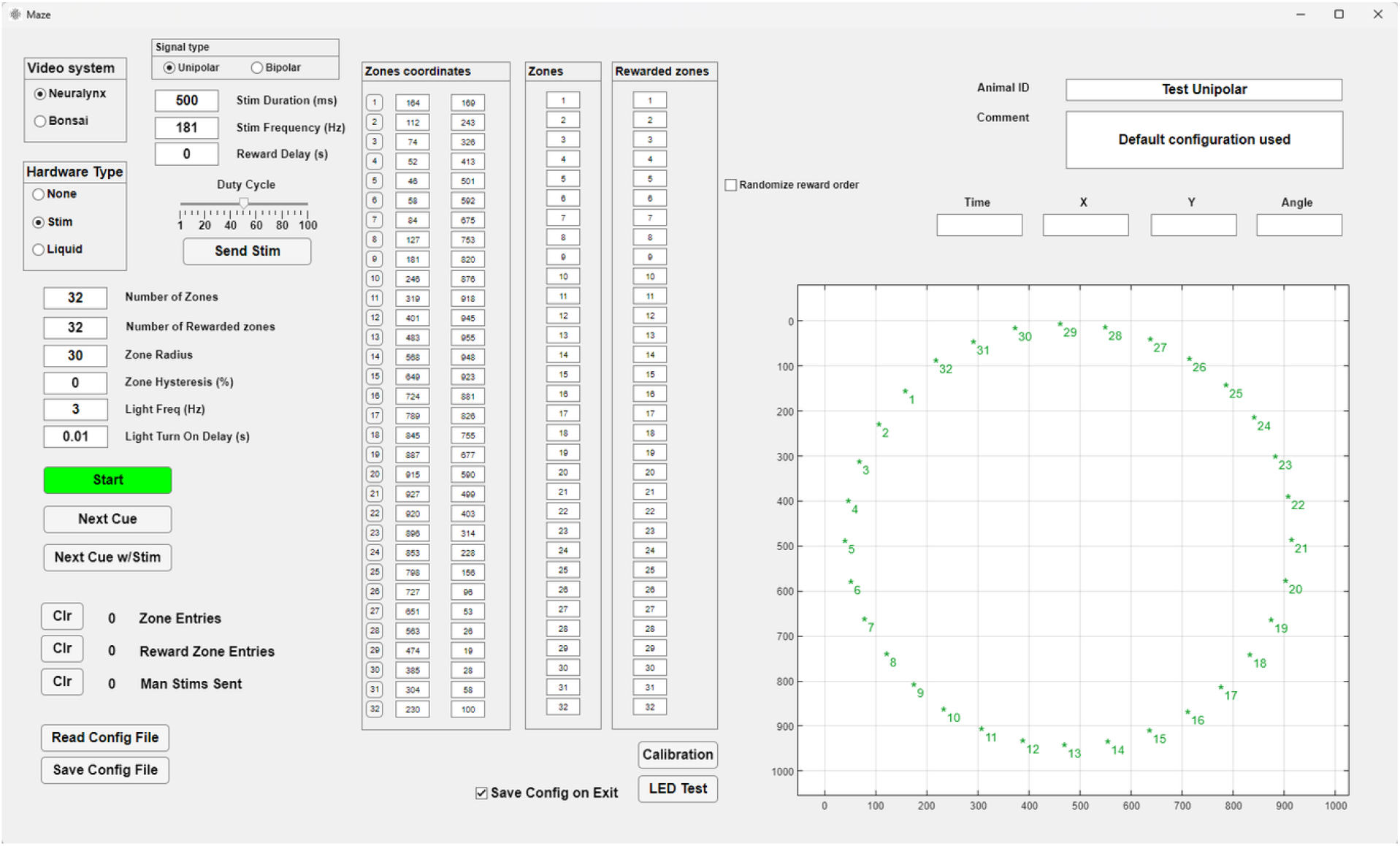
‘Maze’ GUI. Top left: Control of electronic hardware including reward type, stimulation parameters and valve control. Middle left and center: Set the characteristics of zones of interest, including number of zones, zone location, selection of ‘Tracked zones’ and ‘Rewarded zones’, and randomization of rewarded zones. Middle bottom: LED test and auto Calibration controls. Bottom left: Counters and Configuration controls. Top right: Animal ID and Comment boxes. Middle and bottom right: Real-time position tracking.

##### Control of electronic hardware

Users can control all electronic hardware through the GUI. The control for the valves becomes available when users select Liquid reward. If users select Brain stimulation, the panel with the stimulation settings becomes visible and editable, along with a counter for manual (user-triggered) and animal-triggered stimulation. The settings that can be modified include duration, frequency, pulse profile, and delay after entry to the rewarded zone (this feature requires the animal to remain in the reward zone for the experimenter specified delay in order for the animal to obtain a reward). In the ‘Zone coordinates’ panel, buttons corresponding to each zone number allow you to turn the LEDs on and off. When turned on they will become active when the zone is the next/active in the sequence. The LED can be set to blink (or not) and the blink rate can be specified. All changes to the LED status are recorded and timestamped in the acquisition system via TTL sent from the I/O board.

##### Set zone characteristics

The software is designed around spatial zones that are used to record maze events (e.g., when the animal crosses a particular region of the maze) or trigger rewards (e.g., when the animal reaches a specific location). The GUI allows users to set the characteristics of zones of interest. Users can adjust the number of zones to be tracked, and each time an animal enters or exits a zone, the zone number (1-32) is sent via TTL in binary format to the neural signal recording system via the USB-DIO-48 board where the event is precisely timestamped. The zone radius and hysteresis can also be adjusted. Zone hysteresis temporarily increases the zone radius when an animal enters a zone, reducing the likelihood of detecting slight head movements as false zone exits. When the hysteresis option is used, once the animal leaves the zone or the reward is delivered, the zone radius returns to its previous value.

The ‘Zones coordinates’ panel displays the numbers and camera pixel coordinates of up to 32 possible zones, as well as LED control buttons. To determine the location of each zone, users have two options: they can manually enter the xy pixel-coordinates into the input boxes, or they can use the automatic calibration system. With the latter option, the system detects the location of each zone-associated LED using either Cheetah or Bonsai. The LEDs are detected sequentially as they light up from 1 to 32, and the system automatically places a zone at each detected location. The IDs for the zones of interest can be entered in the ‘Tracked Zones’ panel. The ‘Tracked Zones’ are indicated in green font to differentiate them from the inactive zones shown in black. Associating specific LEDs with specific zones allows the user to illuminate those LEDs as cues for the animal, either on every trial or only on user defined trials. Additionally, cue lights can be positioned out of the view of the animal so that the experimenter can use them to ensure the maze software is tracking the animals sequential progress through the maze accurately, or the LEDs can be turned off and not used at all.

##### Zone-triggered rewards

The GUI offers the functionality of zone-triggered rewards. The number of rewarded zones must match the number of zone IDs entered under the ‘Rewarded Zones’ panel in the middle section, and zones must also be included in the ‘Tracked Zones’. Only one zone can be set to be rewarded at one point in time (i.e., until that zone is reached by the animal, or the animals is advanced to the next reward zone manually by the experimenter, a subsequent reward zone – if specified – cannot become active). Upon animal or manual zone triggering, the next zone in the ‘Rewarded Zone’ panel will become active. The currently rewarded zone is encircled in blue in the ‘Animal tracking’ panel. Reward zones can be repeated and can be set up to become active sequentially (if more than one reward zone is used). Randomization of the order in which these zones are selected is also an option. These lists can contain mixtures of rewarded and unrewarded zones (i.e., some zones can be used to track and timestamp the animals progress through key maze segments and others can be timestamped and also paired with an automatic reward if the reward requirements for that zone are met).

Users can select the type of reward the animal receives when it enters one of the active rewarded zones (up to 32 zones can be active for an experiment) through the ‘Reward Type’ selection buttons.

##### Real-time tracking of animal’s position

The GUI displays real-time tracking of the animal’s position in relation to user-defined maze zones of interest. In the bottom right section of the GUI, the ‘Animal tracking’ panel takes up most of the available space. The tracker displays the location of the animal as a red dot on the axes and provides numerical values for the ‘X’ and ‘Y’ coordinates. If a two-color (e.g., LED) tracking system is set up, and HD tracking is enabled in either Cheetah or Bonsai, the angle of the animal’s head is also displayed in the ‘Angle’ text box. Otherwise, the heading is recorded/displayed as 0 degrees. The head direction is visualized as a blue tangent line that shows the orientation of the head.

##### Animal ID and Comment

The right section of the GUI contains the Animal ID and Comment text boxes. The contents of these boxes are saved as a variable in the settings file when a session is ended or when the ‘Save Config File’ button is pressed. This allows the complete configuration of all parameters to be loaded from the same file and stored as a record of the experiment details for that session.

##### Settings file

All the configuration variables are saved in settings files created and loaded in the program. This can be used to quickly store and load different experimental protocols to ensure consistency across days. The settings file includes the zone locations, reward variable, animal’s ID and notes from the loaded session. The settings files are stored in ‘. mat’ format and contain all the settings as variables that can be conveniently accessed with MATLAB© and read in with analysis code without the need to run the Maze software.

#### Stim Trigger

‘Stim Trigger’ provides a simplified GUI for controlling brain stimulation parameters and triggering methods, along with manual stimulation options and a counter to keep track of the number of delivered stimulations without the need of the full featured maze software **(Fig. 3)**.

**Figure 3:**
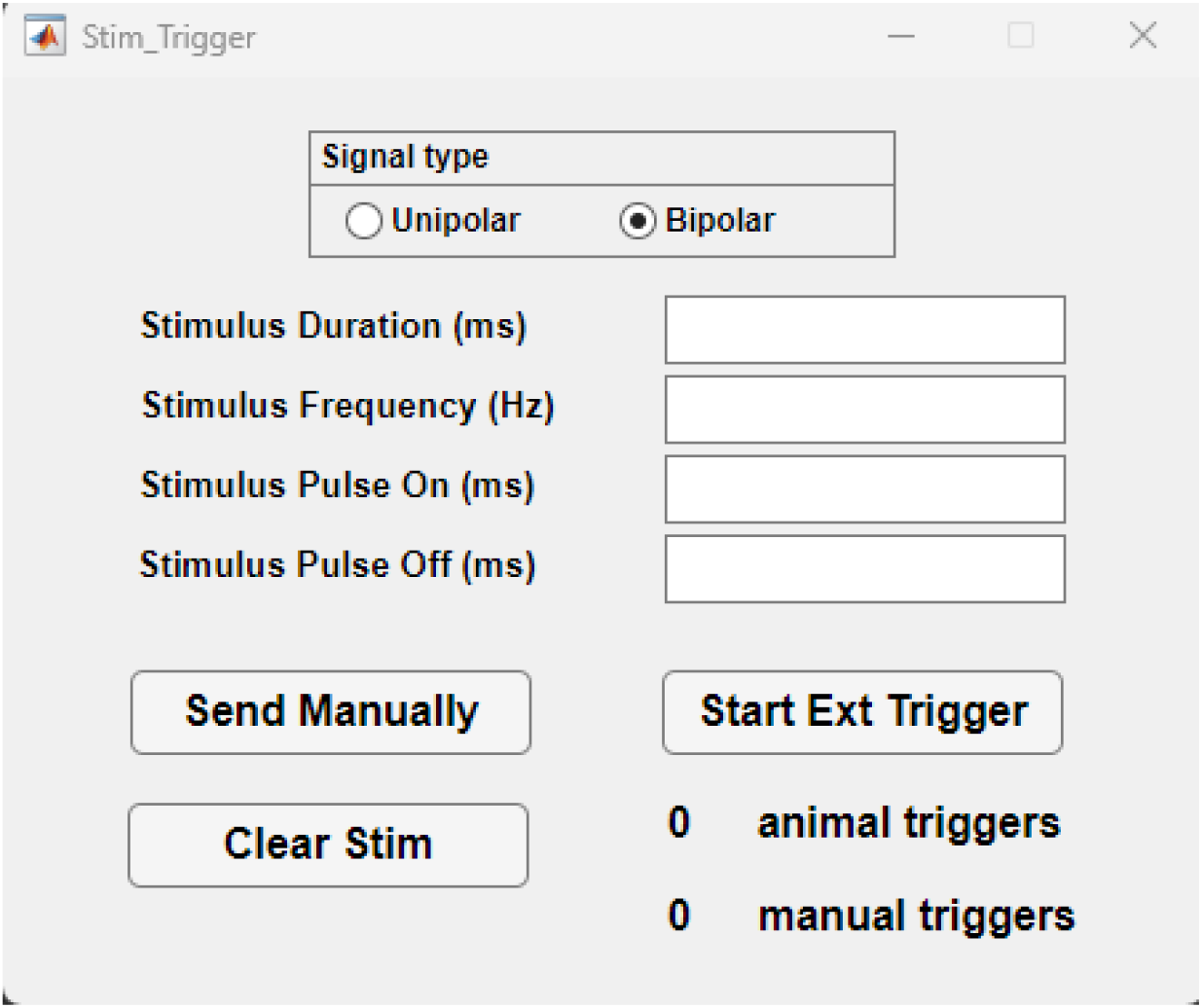
‘Stim Trigger’ software GUI. Top: Stimulus setting input boxes. Middle: “Send Manually” triggers stimulation with each mouse click, “Start Ext Trigger” turns on the automatic trigger, “Clear stim” resets the stimulation counters (right), Bottom: Radio Buttons to select the timing of the automatic trigger on the rising edge or the falling edge of the input TTL from the external triggering source.

Its main purpose is to offer an easy-to-use platform for researchers to control brain stimulation and integrate it with other behavioral equipment or software. The software provides control over electrical or optogenetic brain stimulation, allowing for both manual triggering by the experimenter and automatic triggering by the animal. This stimulation is converted into a bipolar stimulus by the stimulus isolator. Alternatively, a unipolar TTL signal can be utilized to regulate a constant or pulsed laser output for optogenetics. The former method is used alongside a nose poke that operates with a +5V TTL. In addition, a Virtual Reality maze is utilized, incorporating tablets and an Arduino, to deliver electrical brain stimulation rewards when the animal occupies a specific location in the virtual environment (Anon, 2016; Anon, n.d.). Manually triggered stimulation can also be used to shape behavior, such as training animals to approach the nose poke. The software allows users to set the stimulation parameters through the GUI and uses the data acquisition device to trigger the stimulus isolator. As explained above, the duration and frequency determine the main characteristics of the unipolar signal and the duty cycle represents the ratio of time the stimulus is on compared to the time the stimulus is off.

#### Bonsai

Bonsai is a powerful open-source software designed for processing heterogeneous streams of data (https://bonsai-rx.org). It is particularly well-suited for real-time video analysis due to its advanced features and flexibility. In our case, we used LEDs mounted on the recording array to determine the animal’s location and then transferred the location via a UDP port to MATLAB **(Fig. 4)**. Additionally, Bonsai also allows for use of visual methods to determine the animal’s position that do not rely on LEDs, as long as the animal is distinguishable from the background.

**Figure 4:**
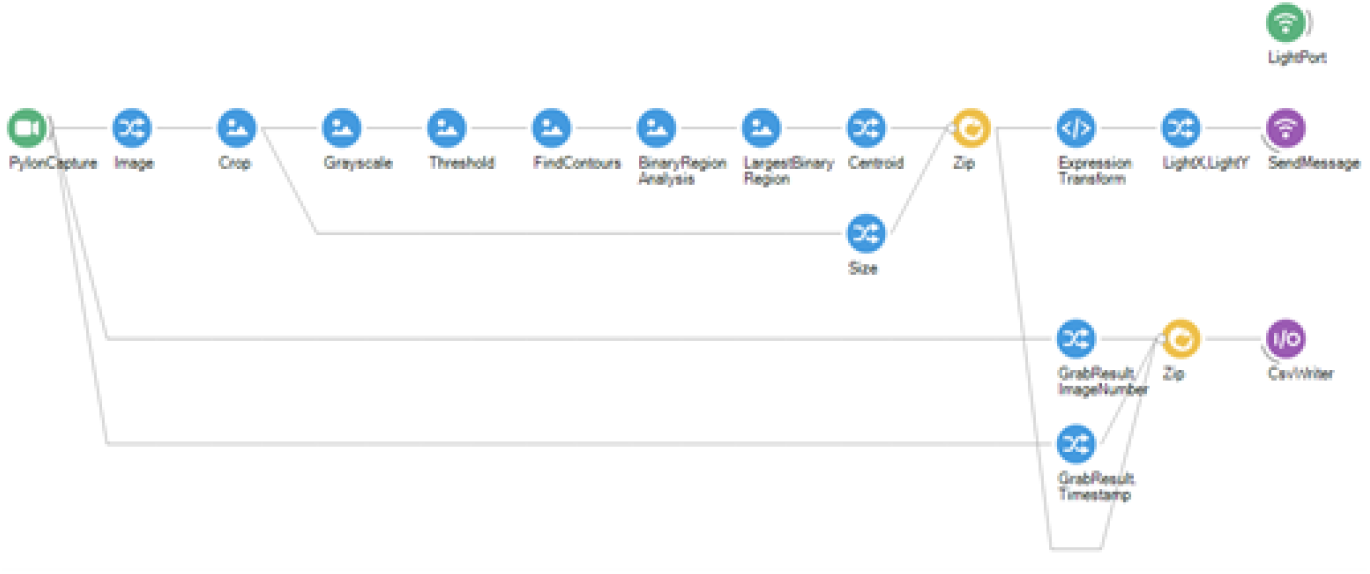
Bonsai workflow for animal’s location extraction and transfer to Maze. Each graphical element represents a function within the data processing pipeline. From left to right, the live video is processed to extract and transmit the animal’s location to both Maze and a .csv file.

The animal’s location is extracted using various functions provided by Bonsai, such as Bonsai.Vision.Threshold, Bonsai.Vision.FindContours, Bonsai.Vision.BinaryRegionAnalysis, Bonsai.Vision.LargestBinaryRegion, and MemberSelector.Centroid. To standardize for different frame sizes the location of the animal is normalized using ‘Size’ from the Video Camera frame. The information extracted by this pipeline is sent using Bonsai.Osc.SendMessage using the UDP port created via Bonsai.Osc.CreateUdpClient.

The threshold settings must be changed to use Bonsai for automatic calibration using the LEDs in the platform. In the presented workflow, we create a .csv file to save the number of frames and timestamps. The location of the animal can also be saved in this file to have a log for the timestamped animal’s location.

## Results

To demonstrate the potential of our platform we will describe a number of applications for this software package across three laboratories and four institutions. The first will be of behavioral data collected from a ‘spatial reorientation’ task that requires mice to use distal cues to get reoriented in space while running down a linear track (Stimmell et al., 2019). Furthermore, we would like to highlight additional publications that have utilized this platform. These include three publications employing a sequence task in rats (Bower et al., 2005; Euston and McNaughton, 2006; Euston et al., 2007), one publication utilizing a task that requires rats to navigate towards a randomly selected cue light out of 32 possible options (Wilber et al., 2014; Wilber et al., 2017) and one using both (Wilber et al., 2017). It is worth noting that all of these publications, involving both the sequence task and the random lights task, incorporated a pre-training phase training rats to shuttle between the ends of a linear track. Additionally, we present an instance where the ‘Stim Trigger’ software was combined with a virtual maze to conduct experiments with mice (Cushing et al., 2020). We also include an example with the Object-Place Paired Associate Task (OPPA) (Sanchez et al., 2019). Finally, we describe its implementation on an unpublished tasks, the Map to Action Transformation (MAT) task.

First, we will elaborate on the application of this software platform for the spatial reorientation task, which was previously described, and has been used for both rats and mice (Rosenzweig et al., 2003; Stimmell et al., 2019). Briefly, the animal must travel back and forth over a linear track with an unmarked rewarded location. The starting point within the linear track is randomized, thus the distance to the rewarded location changes across trials. The animal is moved to a new starting location after each trial while in an enclosed start box where the animal receives a water reward for running to the end of the track and back into the start box.

Prior to running the actual task, and after a week of recovery after medial forebrain bundle electrode implantation, mice were placed in a custom box with a nose poke port. They were trained to approach and poke their nose into port, which triggered a brain stimulation reinforcement. ‘Stim Trigger’ controlled the characteristics of the brain stimulation like duration and frequency for each beam break. The other parameters like current and electrode wire combinations were configured on the stimulus isolator. Adjustments to these were made to achieve the highest response rate over a week. The optimal configuration was used as reinforcer for the entire length of the experiment.

To run the task, the researchers used 3 zone markers: The configuration of the zones was the following: 3 tracked zones, one to start the trial (i.e., exit from the start box), one rewarded zone and one to mark the arrival at the end of the track. As the actual starting point varied in each trial by design, a manual zone skip feature was employed, triggered by a handheld device specifically designed for advancing presentation slides, to mark the beginning of each trial. The virtual location of the ‘Start zone’ was set so the animal could not reach it, and the trial start was marked by pressing the ‘Next Cue’ button using a handheld presentation clicker. Then, the ‘Rewarded Zone’ became active and the animal could trigger the reward, which in this case was electrical stimulation of the medial forebrain bundle. For this application, they used the delay feature for the ‘Rewarded Zone’ so that the animal must remain in the reward zone for a delay period in order to trigger the brain stimulation reward. If the animal advances through the reward zone, or triggers the reward zone, then the next non-rewarded zone, called the ‘End Zone’, becomes active. After reaching the ‘End Zone’, the animal returns to the start box and consumes a water reward while the track is moved to the next randomly selected start location. Thus, at the end of the session the data for the start of the outward trajectory, entry into and exit from the reward zone, reward timestamp if any, and the time the rodent arrived at the end of the track can be extracted.

A post-processing script extracts the animal location from the video file and then calculates the velocity for each trial fixed with respect to the position relative to the reward zone. The velocity profile is visually displayed, enabling the user to identify and exclude trials with issues, such as instances of lost tracking that are too extensive to allow for accurate position estimation by the software. This exclusion process can be performed during visualization of the current session. This analysis code is available upon request.

The results showed that 3xTg-AD female mice performed significantly worse than non-Tg age-matched controls for the 1.5 and 2s delays, but not for other delays (Stimmell et al., 2019). The 6-month non-Tg female mice were able to identify the location of the reward zone and slowed down in preparation for stopping in the zone, whereas the 6-month 3xTg-AD females did not. Overall, the study showed that 6-month 3xTg-AD female mice were impaired at spatial reorientation compared to non-Tg mice.

The virtual version of the task is identical except that the animal’s virtual location is restarted automatically to the next starting location from the end of the track (Cushing et al., 2020). During the virtual task, the animal is placed on a tablet surface coated with mineral oil to facilitate smooth movement of its paws. The animal’s head is fixed to ensure that as it walks, the tablet detects the movement of its paws, thereby navigating the animal through the virtual environment, which is projected on the floor and 3 surrounding wall tablets. Tablet holders are 3D printed with a low-cost resin-based 3D printer and the design files for these tablet holders are available on request. The East, North and West tablets display the rest of the virtual room, which moves along with the animal locomotion. The virtual maze software (Anon, 2016) was paired with ‘Stim Trigger’ which controlled the issuance of brain stimulation as described above.

Our next detailed example includes *in vivo* electrophysiological data. Rats were first trained on an alternation training task, where they learned to shuttle back and forth along a linear track created by walls that connect two opposite zones on the circular platform. Rats receive a brain stimulation that is delivered in at both ends of the linear track. To indicate the alternate goal location for the animal, an LED would blink at each end, serving as a cue for the experimenter. Following the alternation training, rats underwent training on a ‘random lights task’, where sequences of up to 900 elements were drawn randomly with replacement from the 32 light/reward zones.

Subsequently, rats were trained on the complex spatial sequence task, which involves rats learning to navigate to unmarked locations fixed in space in a specific sequence. Landmarks are distributed around the room for spatial orientation. We used a sequence (1-2-3-4-1-2-3-5-).; **(Fig. 5)**; **(Vid. 1)**) that had a repeating path segment (1-2-3) followed by one of two distinct actions. Specifically, the rat learned in context 5-1-2-3, to go to 4 for reward, while in context 4-1-2-3 the rat must go to 5 **(Fig. 5)**. Thus, each action belongs to two spatial contexts, so navigation to zone 4 or 5 requires a map-to-action transformation. This emulates the common spatial memory problem one encounters when driving through an intersection and remembering the appropriate action given the current route and goals (e.g., turn left to a favorite restaurant versus right to home). Sets of 3 unguided (‘memory’) runs through the complete 1-2-3-4-1-2-3-5 sequence were interleaved with sets of 3 ‘cued’ runs in which a light at each goal led the rat through the sequence. During memory runs, following an error, a light cue directs the rat to the next zone in the sequence. This complex spatial task, while likely engaging the hippocampus, hippocampal activity alone (e.g., splitter cells) is insufficient to predict the rat’s action (Bower et al., 2005). Note, because the sequence makes use of 5 zones selected from 32 evenly spaced zones distributed around the perimeter of the platform, alternate sequences which match the distance traveled for various elements of the sequence can be created or new sequences can be created by flipping and rotating the sequence to make a novel sequence (we have made use of both of these options in our experiments). The data presented here is the activity of two place cells recorded from the rat’s dorsal hippocampus while it navigated the 8-item sequence on circular platform. The cells have distinct firing patterns focused in a specific location on the platform **(Fig. 6)**. This method allowed to train the animal by reinforcing their behavior via brain stimulation or food reward. Rats have been trained under both conditions on the three tasks described in this section. Additionally, this method allowed the researchers to detect the animal’s precise location during behavior, and record the neuronal activity with high temporal resolution, using the recording system.

**Figure 5:**
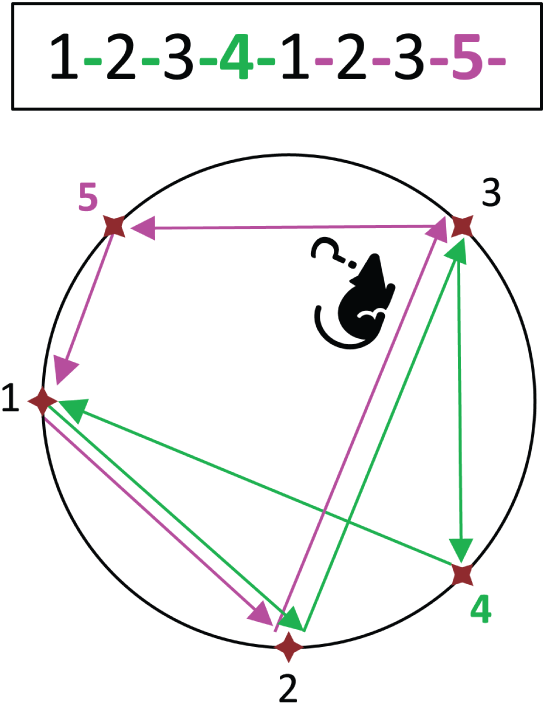
Complex Spatial Sequence Task. Schematic for the complex spatial sequence task. The rat always starts at zone 5 and continues to zone 1-2-3-4-1-2-3-5-.

**Figure 6:**
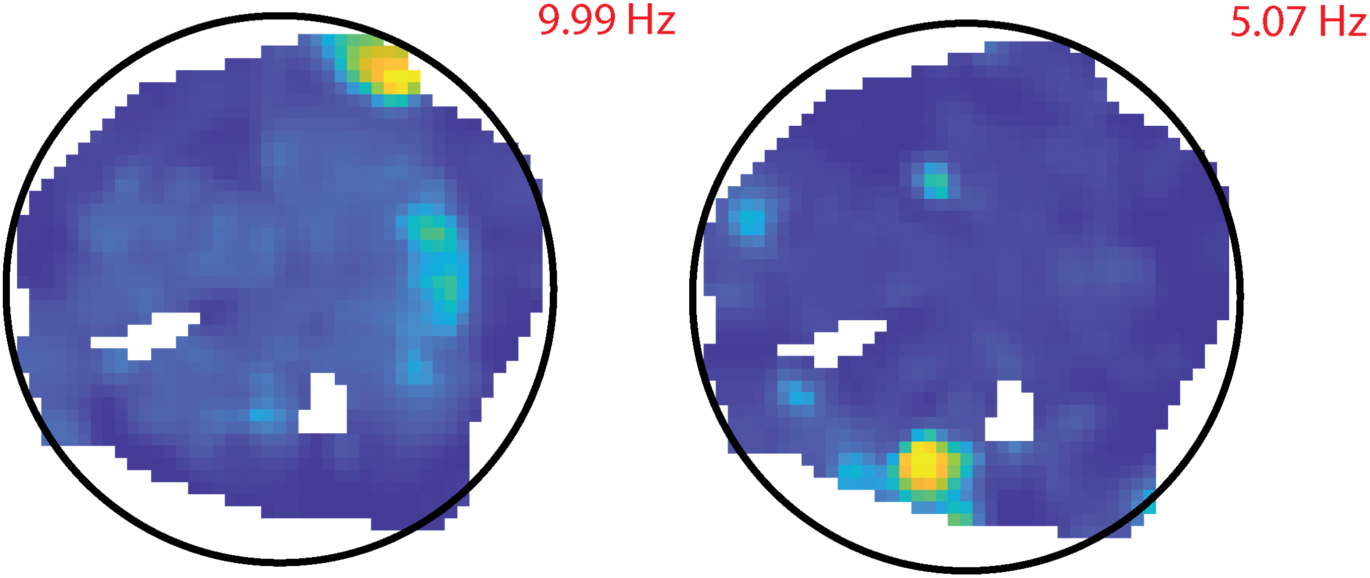
Two plots showing the activity of two hippocampal place cells as a heat plot of time-adjusted firing rate, using an evenly spaced color-map with max rate indicated in red. The firing frequency is also shown. Only areas with a high enough occupancy during the task are represented in the figures.

**VIDEO 1.** ‘Maze’ software configuration for Sequence task. At the beginning of the video, a configuration file containing all the necessary settings for Sequence task is loaded. Then, the animal is connected and electrophysiology recording and live location data are initiated. Subsequently the ‘Maze’ program is initiated and the location of the animal is displayed in the bottom right corner of the software window. In the second part of the video, a rat completes the Sequence task by navigating between the rewarded zones. The path that the animal must travel from memory in order to obtain rewards at each zone is overlaid on the recording for easy visualization.

The next example of the use of our platform is with the Object-Place Paired Associate Task (OPPA). Previously described in (Sanchez et al., 2019). In brief, rats are trained to travel to the end of a two-arm maze where they have to displace a specific object to obtain a reward out of two possible options. The animal must then navigate back to the center of the maze and go to the other arm. The goal of the current project is to analyze the local field potential on each maze arm, excluding the choice point where the objects are located. To do this, the researchers are using our Maze software to get accurate timestamps of the location of the animal, as well as entry in zones of interest. They use 7 zones, one for each platform at the end of the arms, two for the maze arms, one for the central platform of the maze and two zones outside of the maze to timestamp the manual reward delivery **(Fig. 7)**. To obtain the location of the animal, they use Bonsai which sends the location data via UDP, as well as head direction (angle) to the maze software. Each zone entry is sent to the recording system as a TTL to obtain the precise timestamps of the maze events for post processing.

**Figure 7:**
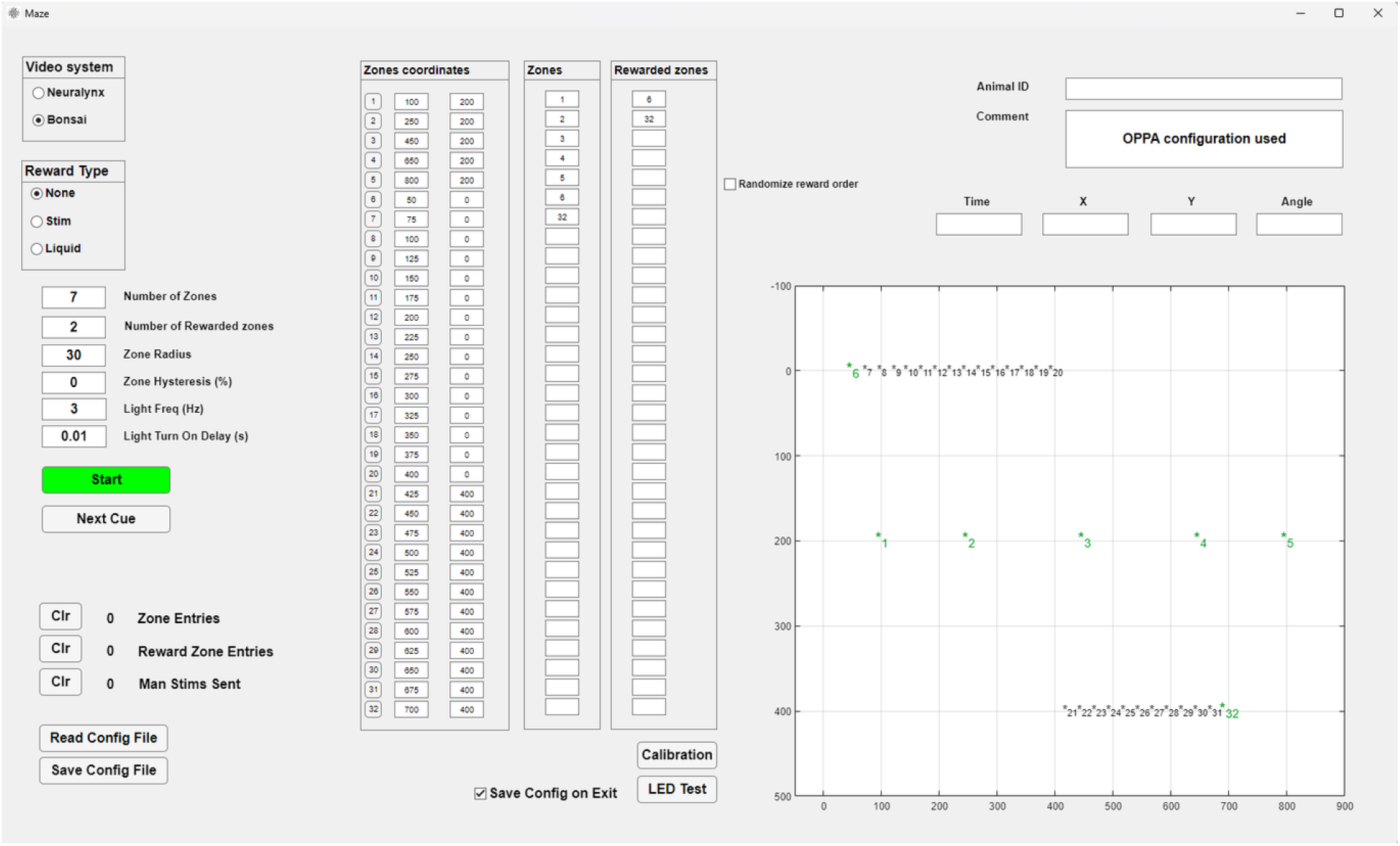
Maze GUI for OPPA task. Top left: Control of electronic hardware including reward type: None as manual food reward is used. Middle left and center: 5 zones of interest, with allocated coordinates for zone location. Unused zones are listed above and below the zones that are positioned on the maze for visualization. Bottom left: Counters and Configuration controls. Top right: Animal ID and Comment boxes. Middle and bottom right: Real-time position tracking with zones.

The final example is the Map-To-Action Transformation (MAT) Task. Rats first learn to navigate to the reward location (R in **Fig. 8** Left) from a randomly selected set of 7 start locations (S1-7) using distal cues. The maze is surrounded by landmarks for orientation. Accurate navigation results in food or brain stimulation reward delivered at the reward location. After achieving allocentric condition criterion (over 80% correct trials for at least 3 out of 4 sessions), the transformation condition begins. In this condition, the rat is held at each start location in a translucent box for 10s. This is done for Allocentric and Egocentric conditions to produce parallel data sets for each task. Next, an opaque box is placed over the rat while curtains are pulled obscuring distal cues (**Fig. 8** *Middle*). The box is then removed, and the rat must now navigate to R while the distal cues are obscured. The hypothesis is that in this condition the animals must transform the allocentric location of R into an egocentric action sequence to reach the reward.

**Figure 8:**
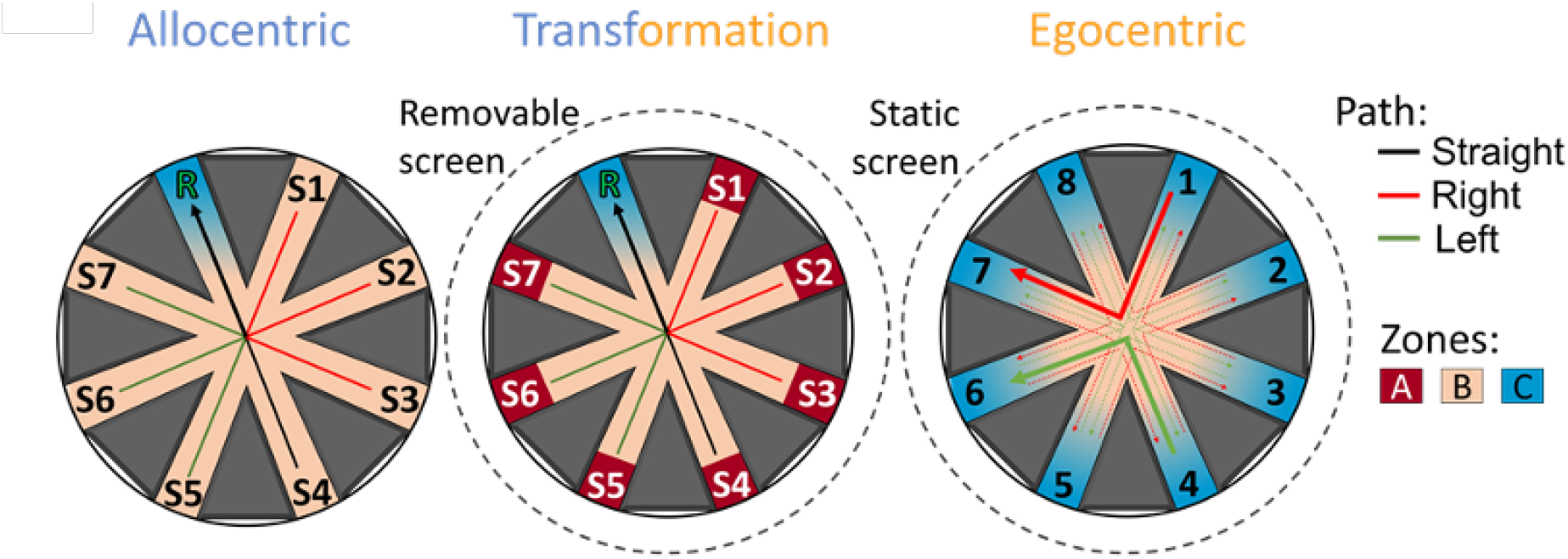
Map-To-Action (MAT) task. Layout for the allocentric (Left) transformation (Middle) and ego-centric conditions (Right) of the MAT task.

After achieving the same criterion for the map-to-action transformation condition, the egocentric paradigm begins where the egocentric relationship between the start location and the rewarded location is maintained (e.g., 4-6). However, these locations are randomized for each trial while the distal cues are obscured by the curtains. There are two variations, right turn (**Fig. 8** Right Red and left turn, Green). Each rat runs both the right turn and left turn variation separately, and the order is counterbalanced across rats. In this condition, the rat needs to develop a self-centered motor strategy to find the reward and cannot rely on any external cues which are unavailable (distal) or changed every trial (proximal).

Spatial zones are positioned in ‘Maze’ software order to timestamp zone entry for each of the 8 arms as well as the center of the maze. Some animals are trained with food reward, while others with electrical stimulation of the medial forebrain bundle as the reinforcer. For the animals that receive brain stimulation reinforcement, the delivery of the brain stimulation pulse train via TTL to the recording system is also timestamped. For all training phases, a non-rewarded zone is positioned in the center of the maze to timestamp entry into the center of the maze. In the case of the allocentric and transformation tasks, the rewarded location remains consistent throughout the session. Two rewarded zones are designated, one located within the correct zone and the other positioned outside of the maze. This ensures that upon zone entry, the reward is delivered only once until the experimenter utilizes the ‘Next Cue’ button to reactivate the rewarded zone on the maze. The zone that is positioned off the maze is also used to signal the end of an incorrect trial.

Due to the continuously changing rewarded location for the egocentric task, the ‘Stim Trigger’ software is used to deliver and timestamp the reinforcement delivery for correct trials. Furthermore, manually triggered TTLs can mark events that are not location-dependent, such as the animal placement into boxes. To do this, a wireless relay switch activated by a wireless radio frequency remote control can be used. This relay system is connected to the Neuralynx TTL acquisition board. Four control rats have successfully completed all three paradigms of the task. Training for each phase takes a similar number of sessions, between 19 and 23.

## Discussion

In spatial cognition research, it is essential to have the flexibility to quickly change the experimental setup without having to overhaul the software and hardware. To address this need, we present an inexpensive behavioral platform that can be paired with open-source or commercial software to create a variety of experimental setups. Our platform enables the generation of novel spatial navigation tasks, using positive reinforcement like brain stimulation, liquid, or food reward. We are presently using this platform in our laboratory to collect data using eight different mazes/experimental paradigms and there are countless more experiments that can be designed and run using this platform.

Our software includes two highly flexible programs. The Maze program can use up to 32 user defined zones and deliver rewards automatically upon zone entry or track progress through the zones. This program is highly synergistic with electrophysiology acquisition systems like Neuralynx and OpenEphys, as it timestamps zone entries and other behaviors using TTL signals triggered by the Maze program and delivered to Digital Lynx SX or OpenEphys Acquisition Board. The second program, ‘Stim Trigger’, can control brain stimulation using any type of equipment that can be paired with an Arduino board, such as a nose-poke and a virtual reality maze.

Our platform distinguishes itself from other commercial and open source software options for animal behavioral tracking by offering more than just tracking capabilities. For example, EthoVision is a commercially available system focused on video tracking, our platform goes beyond tracking to provide a versatile and flexible environment for experimental design. Additionally, the modular design of our platform allows researchers to incorporate a wide range of hardware and software components into their experiments, making it highly adaptable while keeping costs low.

In contrast to other commercial solutions like ANY-maze, our platform is freely available, either in a compiled version or with MATLAB compatibility, and can be edited and customized according to researchers’ specific needs. ANY-maze, although versatile, is not free and lacks intrinsic ability to synchronize with other software platforms like Python or MATLAB.

Finally, our platform seamlessly integrates with Bonsai-RX and DeepLabCut, enhancing its capabilities even further. With Bonsai-RX, researchers can perform real-time behavioral analysis and closed-loop experiments, while DeepLabCut’s deep learning algorithms enable accurate and automated tracking of key points on the subject’s body, allowing for precise analysis of movement and behavior. The synergy between our platform and these software solutions facilitates the implementation of complex experimental setups, incorporating multiple data streams such as video tracking, electrophysiology, and optogenetics, offering researchers extensive possibilities for data analysis and visualization.

The purpose of sharing these programs is twofold: First to share the software as is, and second, to get feedback from the community outside of our groups on ways to improve and expand our software package. We have focused on making our software compatible with open-source software and hardware like Bonsai and OpenEphys. A potential area for expansion is with another cutting-edge open-source technique, the UCLA Miniscope (Ghosh et al., 2011). It can be paired with our ‘Maze’ software using the Miniscope V4 - Data Acquisition System to synchronize Neuralynx or Bonsai (with a blinking LED for synchronization) video acquisition with the physiological data and timestamp location information. We seek to continue developing these tools and collaborating with other researchers to increase the possible applications of our work.

## Limitations

Our software has limitations when it comes to compatibility with all experimental designs. For instance, the current version only allows for the delivery of rewards upon entry to a zone or zones sequentially, but two rewarded zones cannot be available simultaneously, as we have not yet encountered a need for this functionality.

While our software has been effectively used in both rats and mice, including several publications involving mice, it is important to note that the physical maze platform described in the methods section was primarily designed with larger rodents, such as rats, in mind. To address this, we propose the creation of scaled-down versions of the platform specifically tailored for smaller rodents. These scaled-down versions would function in a similar manner to the original platform, with the location and size of the zones adjusted proportionally based on the size of the camera frame. No modification of the software is needed to make use of a scaled down platform.

We also plan to address compatibility issues. While the software and hardware are compatible with open-source software and hardware such as Bonsai and OpenEphys, we recognize that MATLAB © is not freely available. Therefore, we aim to transition to other programming languages such as Python in the future. In the meantime, MATLAB can be used to compile the program (and we have made available a compiled version of the current program), so MATLAB is only needed when making edits to the program. While we acknowledge that a single platform may not be able to cover all experimental designs, the software’s flexible structure enables us and other users to create custom versions quickly.

## Data availability

The presented code, except analysis code, is available in our data repository in OSF, link:https://osf.io/svtzr/. The list of components to build the platform is available on Google Sheets, link: https://docs.google.com/spreadsheets/d/1GMOYG8HO3yyJB4AWzgjLCo-qlfUHs2DzC6-nCg6M2rA/edit?usp=sharing

## Acknowledgements

N\A

## Notes

**Conflict of Interest** Authors report no conflict of interest

https://osf.io/svtzr/

### Competing Interest Statement

The authors have declared no competing interest.

### Summary of Updates

Figure 5 revised to increase image quality.

